# An Expanded Class of Histidine-Accepting Viral tRNA-like Structures

**DOI:** 10.1101/2020.12.02.408831

**Authors:** Conner J. Langeberg, Madeline E. Sherlock, Andrea MacFadden, Jeffrey S. Kieft

## Abstract

Structured RNA elements are common in the genomes of RNA viruses, often playing critical roles during viral infection. Some RNA elements use forms of tRNA mimicry, but the diverse ways this mimicry can be achieved are poorly understood. Histidine-accepting tRNA-like structures (TLS^His^) are examples found at the 3′ termini of some positive-sense single-stranded RNA (+ssRNA) viruses where they interact with several host proteins, induce histidylation of the RNA genome, and facilitate several processes important for infection, to include replication. As only five TLS^His^ examples had been reported, we explored the possible larger phylogenetic distribution and diversity of this TLS class using bioinformatic approaches. We identified many new examples of TLS^His^, yielding a rigorous consensus sequence and secondary structure model that we validated by chemical probing of representative TLS^His^ RNAs. We confirmed new examples as authentic TLS^His^ by demonstrating their ability to be histidylated *in vitro*, then used mutational analyses to verify a tertiary interaction that is likely analogous to the D- and T-loop interaction found in canonical tRNAs. These results expand our understanding of how diverse RNA sequences achieve tRNA-like structures and functions in the context of viral RNA genomes and lay the groundwork for high-resolution structural studies of tRNA mimicry by histidine-accepting TLSs.

## INTRODUCTION

Viruses are obligate cellular parasites that must subvert and co-opt host cellular machinery to proliferate. In single-stranded RNA (ssRNA) viruses, structured regions within the viral genomic RNA can directly manipulate cellular machinery, a ubiquitous part of many viruses’ overall infection strategy. Such RNA structures can affect pathways and processes as diverse as translation, replication, packaging, viral RNA stability, and immune evasion (Pathak et al. 2011; Tuplin 2015; Garcia-Blanco et al. 2016; Hogg 2016; Jaafar and Kieft 2019). RNA structural elements frequently act to coordinate the switch between different processes occurring on the genomic RNA, often using changes in conformation to do so (Gamarnik and Andino 1998). Understanding the structural diversity and distribution of diverse viral RNA elements is essential to define their mechanisms of action within the overall viral infection process. More generally, exploration of viral RNA structure helps us understand roles for RNA structure in diverse cellular processes by revealing the fundamental rules for how RNAs fold to drive function. In particular, because ssRNA viruses evolve relatively rapidly, exploring conservation of sequence and structure of an RNA class within different viruses reveals how dissimilar RNA sequences achieve a similar structure and function (Mans et al. 1991; Roth and Breaker 2009; Webb et al. 2009; Perreault et al. 2011; Steckelberg et al. 2018; Pisareva et al. 2018; Jones et al. 2020).

A class of structured RNAs that play roles in infection are the tRNA-like structures (TLSs), found at the 3′ terminus of certain positive-sense single-stranded RNA (+ssRNA) viruses (Rietveld et al. 1984; Mans et al. 1991; Dreher 2010). These RNAs mimic the structure of transfer RNAs (tRNAs) to varying degrees (Felden et al. 1994a, Felden et al. 1996; Hammond 2009), and functionally mimic tRNAs in multiple ways. Notably, these RNAs can be aminoacylated on their 3′ adenosine by host cell aminoacyl tRNA synthetases (AARSs) (Pinck et al. 1970; Ijberg and Philipson 1972; Kohl and Hall 1974; Salomon et al. 1976; Joshi et al. 1985; Goodwin and Dreher 1998). Furthermore, they interact with host proteins associated with tRNAs, specifically eukaryotic elongation factor 1A (eEF1A) (Joshi et al. 1986; Zeenko et al. 2002; Hwang et al. 2013; Li et al. 2013), and the CCA-adding enzyme (S. Litvak, L. Tarrago-Litvak 1973; Dreher and Goodwin 1998; Hema et al. 2005). Viral proteins are also known to interact with TLSs, which contain part or all of the viral negative-strand promoter site required for RNA-dependent RNA polymerase (RdRp) binding (Singh and Dreher 1997; Deiman et al. 1998; Osman et al. 2000; Olsthoorn et al. 2004; Yamaji et al. 2006; Rao and Cheng Kao 2015). In addition, in some viruses the TLS is essential for proper packaging of the virion, acting as a nucleation site for capsid assembly (Choi and Rao 2000; Choi et al. 2002). Thus, TLSs are multifunctional RNAs, a feature conferred by their three-dimensional structure.

Studying TLSs may give insight into other types of tRNA mimics proposed to exist in viral and cellular RNAs. Other putative tRNA mimics include the tRNA-miRNA-encoded RNAs (TMERs) in Gammaherpesvirus (Diebel et al. 2015) and those in some 3′ cap-independent translational enhancer (3’ CITE) elements (McCormack et al. 2008; Simon and Miller 2013). Non-viral tRNA mimics include transfer-messenger RNAs (tmRNAs) (Williams and Bartel 1995; Weis et al. 2010) and the MALAT1-derived mascRNA (Wilusz et al. 2008; Sun and Ma 2019; Lu et al. 2020). Notably, while some of these enhance translation, none are known to be aminoacylated, and while some remain within the genome, others are processed out of the primary transcript. TLSs remain intact within the viral genome but must use a pseudoknot to mimic the acceptor stem in a manner that both allows aminoacylation at the 3′ terminus and connectivity with the viral genome upstream of the 5′ end of the TLS.

Three classes of TLSs are known: valine-accepting TLSs (TLS^Val^), tyrosine-accepting TLSs (TLS^Tyr^), and histidine-accepting TLSs (TLS^His^) (Mans et al. 1991; Dreher 2010). Each class is distinct in the identity of the amino acid added to its 3′ end, secondary structure elements, and presumably their higher-order folding, but all contain a pseudoknotted acceptor stem mimic as well as a 3′ CCA added by cellular processing machinery. The TLS^Val^ class most resembles a canonical tRNA structure with discernable acceptor stem, D-arm, T-arm, and anticodon (AC)-arm elements (Pinck et al. 1970; Fukai et al. 2000; Hammond et al. 2009), confirmed by two crystal structures of the turnip yellow mosaic virus TLS (Colussi et al. 2014; Hartwick et al. 2018) Conversely, the TLS^Tyr^ class appears to be the most divergent from tRNAs, with several additional stem-loops present, no clear T-arm or AC-arm, and thus no obvious way to mimic tRNA (Haenni et al. 1982; Dreher and Hall 1988; Felden et al. 1994a). However, a recent cryo-EM structure of the Brome mosaic virus (BMV) TLS^Tyr^ revealed tRNA mimicry is embedded in a more complex fold that may require programmed conformational changes to fully create the tRNA mimicry (Bonilla et al. 2020).

The third class of TLSs, TLS^His^, visually appears to lie between the other two classes in terms of structural similarity to canonical tRNA. The TLS^His^ secondary structure has putative analogs to the AC-arm, T-arm, and acceptor stem, though it lacks an obvious D-arm analog (Ijberg and Philipson 1972; Salomon et al. 1976; Rietveld et al. 1984; Felden et al. 1996). High-resolution structural insight into this class has remained elusive, likely due in part to the structural heterogeneity observed in the prototypical TLS^His^ RNA from the tobacco mosaic virus (TMV) (Hammond et al. 2009). Although structural modeling of the RNA provided insight into the possible three-dimensional fold of the TMV TLS (Rietveld et al. 1984; Felden et al. 1996), this model remains untested. Furthermore, only five TLS^His^ sequences have been identified, which makes analysis of this class challenging compared to the TLS^Val^ and TLS^Tyr^ classes (Dreher 2010; Sherlock et al. 2020; Bonilla et al. 2020). Hence, although the TLS^His^ class represents a novel form of tRNA mimicry that can provide insight into diverse ways this form of mimicry can be achieved and used, it is largely mysterious.

The relative scarcity of sequence and structural information available for the TLS^His^ class motivated us to better understand how primary sequence, secondary structure, and tertiary contacts achieve the proper fold required for their function. Using the few previously reported TLS^His^ sequences, we performed bioinformatic searches based on primary sequence conservation and secondary structure patterns to identify many additional putative TLS^His^ sequences. We used chemical probing to confirm the proposed secondary structures of some new TLS^His^ and used an *in vitro* aminoacylation assay to verify that they are novel, functional, histidine-accepting TLSs. Finally, we interrogated a proposed D-loop mimic, implicating this region in a long-range interaction with the T-loop that is reminiscent of the D-loop/T-loop interaction present in canonical tRNAs. Together, our findings uncover an expanded phylogenetic diversity of the TLS^His^ class and provide insight into how the structural conservation of these RNAs correlates with their tRNA mimicry.

## RESULTS AND DISCUSSION

### Bioinformatic searches reveal additional TLS^His^

Identifying new TLS^His^ examples promised to reveal conserved regions required for achieving the proper structure, likely locations of protein interactions required for function, and variations such as the insertions found in the divergent members of the TLS^Val^ and TLS^Tyr^ (Sherlock et al. 2020; Bonilla et al. 2020). To identify additional putative TLS^His^ RNAs that conform to the previously proposed secondary structure model, we performed homology-based searches using the program Infernal (Nawrocki and Eddy 2013). We generated the initial seed alignment from the four TLS^His^ sequences deposited in the Rfam database (Rfam ID: RF01077) (Kalvari et al. 2018), which we modified to include the entire 3′ end, to include the T-loop and acceptor stem pseudoknot. We bioinformatically identified 158 unique sequences from 36 unique viruses (see Supplemental Files 1 and 2). The difference between the numbers of unique sequences and unique viruses results from the presence of subgenomic RNAs with sequence variations between isolates and strains. Of the 36 viruses containing a putative TLS^His^, 33 belong to the *Tobamovirus* genus, two to the *Tymovirus* genus, and one to the *Furovirus* genus (Fig. 1A, Supplemental File 2). Although the tobamoviruses and furoviruses are both in the *Virgaviridae* family and are closely related based on RdRp sequence, the tobamovirus TLS^His^ are more similar to those from the *Tymovirus* genus in the *Tymoviridae* family. The single TLS identified from *Furovirus* had a high E value of 0.019, making it an unlikely TLS^His^ candidate; rather, it demonstrates features of the TLS^Val^ class and was excluded from further analyses. Interestingly, the tymoviruses mostly contain TLS^Val^ and most examples of TLS^Val^ are within *Tymoviridae* with a few in *Virgaviridae* (Sherlock et al. 2020). While we cannot propose a specific evolutionary history of these viral lineages, this distribution suggests exchange of TLS elements between viruses can occur, likely during coinfections by two distinct viruses.

**Figure 1.**
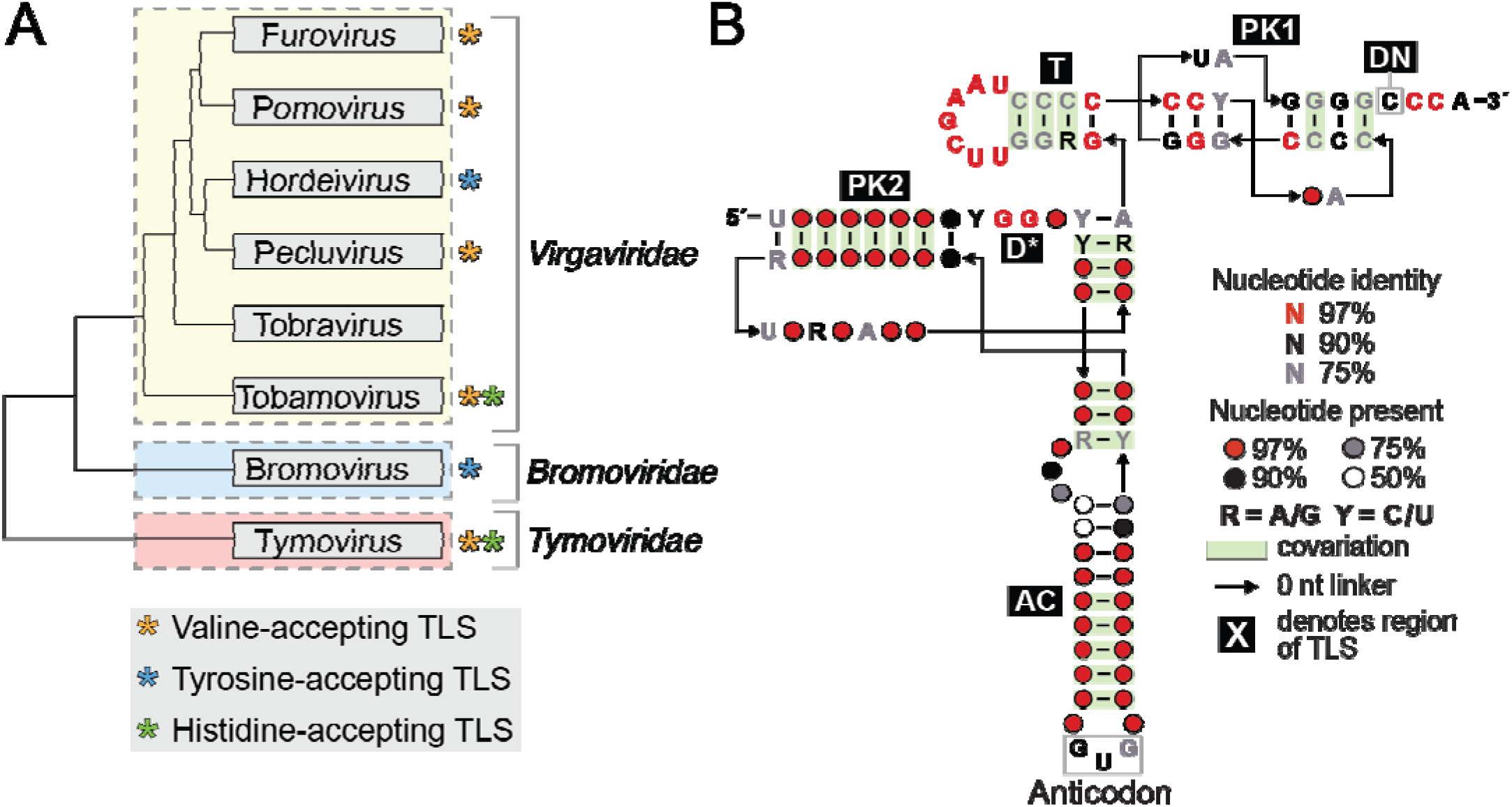
Histidine-accepting tRNA-like structures. **A)** Phylogenetic distribution of tRNA-like structures in several +ssRNA plant virus genera. Tree is based on Met-Hel-RdRp sequence, adapted from (King et al. 2012). Asterisks denote the presence of one or more tRNA-like structure classes in each viral genus. **B)** Consensus sequence and secondary structural model of the 157 identified unique histidine-accepting t like structure sequences. Regions are labeled relative to their homology in a canonical tRNA, if present. PK2: pseudoknot 2 region, D*: putative D-loop analog, AC: anticodon arm, T: T-arm, PK1: pseudoknot 1 NA-region, DN: discriminator nucleotide.

### TLS^His^ sequence conservation and predicted secondary structure patterns

We calculated a consensus sequence and secondary structure model using the R-scape program with the 157 putative unique TLS^His^ sequences as input (Fig. 1B) (Rivas et al. 2016, Rivas et al. 2020). The resultant covariation patterns support the putative secondary structure of the prototypical TMV TLS^His^. Specifically, both pseudoknots and analogs for the T-arm and AC-arm were present in our model and in good agreement with the observed base pairing and covariation.

Interestingly, the degree of sequence conservation is markedly different in different regions of the RNA secondary structure. Specifically, the sequence conservation is high in the region that contains the acceptor stem analog PK1 and the T-arm analog (3′ region). The combined length of the T-arm combined with PK1 is always 11 base pairs, one base pair shorter than in the TLS^Val^ class (Sherlock et al. 2020). Within the T-arm, the T-loop is nearly perfectly conserved in TLS^His^ sequences, containing the 5′- UUCGAAU-3′ sequence common in tRNA T-loops. In fact, the T-loop of the TLS^His^ is more conserved than the T-loop of canonical tRNA^His^ (Westhof and Auffinger 2001). The size and sequence conservation in this region likely reflects how these TLSs are recognized by various proteins, including host HisRS, CCA-adding enzyme and eEF1A, and viral RdRp (Hegg et al. 1990; Singh and Dreher 1997; Deiman et al. 1998; Osman et al. 2000; Zeenko et al. 2002; Olsthoorn et al. 2004; Yamaji et al. 2006; Hwang et al. 2013; Li et al. 2013). Indeed, key bases for recognition by host HisRS are in this region (Crothers et al. 1972; Hou 1997; Rudinger et al. 1997; Tian et al. 2015) and both the CCA-adding enzyme and eEF1A bind to this part of tRNAs (Xiong and Steitz 2004; Nissen et al. 1995).

In contrast to the 3′ region, the region comprising PK2 and the AC-arm analog (5′ region) exhibits substantial base pair covariation but little primary sequence conservation, thus a specific secondary structure is required independent of nucleotide identity. The exceptions are two isolated motifs that are well conserved at the primary sequence level: the histidine AC (GUG) and a GG dinucleotide adjacent to PK2. Finally, we did not find any new TLS^His^ with substantial insertions or deletions as has been seen in both the TLS^Val^ and TLS^Tyr^ classes (Sherlock et al. 2020; Bonilla et al. 2020). Cumulatively, these bioinformatic patterns suggest a likely conserved secondary structure present in histidine-accepting TLSs that matches the proposed structure of the archetypal TMV TLS^His^ and less global variation (insertions or deletions) than is seen in the other classes.

## TLS^His^ adopts a conserved secondary structure

While the high degree of covariation present in all base pairing regions supports a common TLS^His^ secondary structure, experimental interrogation of representative RNAs is critical to test the models. We applied selective 2′ hydroxyl acylation analyzed by primer extension (SHAPE) *in vitro* chemical probing, which queries the conformational flexibility at each nucleotide position in an RNA (Yoon et al. 2011; Kim et al. 2013; Cordero et al. 2014; Kladwang et al. 2014; Lee et al. 2015). Locations in the RNA which undergo a higher degree of motion, typical of an unpaired or otherwise unstructured base, react more readily with the SHAPE reagent *N*-methyl isatoic anhydride (NMIA) than those in base pairs or other interactions that restrict motion. Mapping these relative reactivities onto the proposed structure is then used to validate or adjust the initial model. We applied this method to the putative TLS^His^ from Tobacco Mosaic Virus (TMV), Odontoglossum Ringspot Virus (ORSV), Ribgrass Mosaic Virus (RMV), Cucumber Mottle Virus (CMoV), Maracuja Mosaic Virus (MarMV), Hibiscus Latent Fort Pierce Virus (HLFPV), Zucchini Green Mottle Mosaic Virus (ZGMMV), and Diascia Yellow Mottle Virus (DiaYMV). In all cases, we observed reactivities reflecting the proposed secondary structure of each RNA (Fig. 2, Fig. S1). Specifically, low reactivities were observed in regions proposed to be base-paired, namely both pseudoknot regions, the AC stem, and the T-arm stem. Conversely, most regions predicted to lack canonical base pairing in the model contained elevated levels of reactivity: the AC loop, the linker regions following PK2, the T-loop, and the CCA trinucleotide. Additionally, the internal loop or bulge present in all AC stems was highly reactive, consistent with conformational dynamics in this region. This loop often contains five nucleotides but varies in size (Fig. 2, Fig. S1).

**Figure 2:**
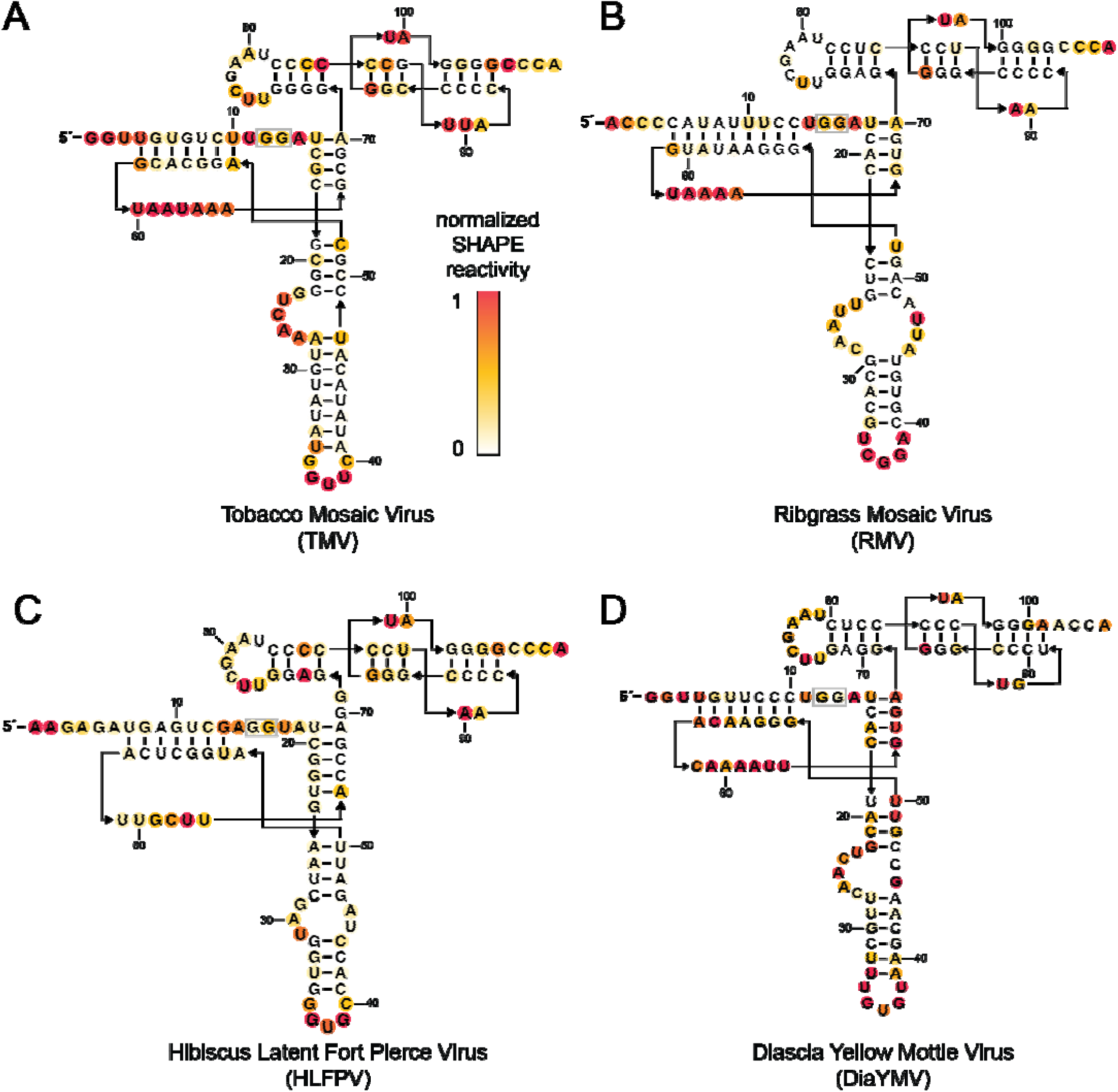
Histidine-accepting tRNA-like structures adopt a conserved secondary structure. Chemical probing of four representative TLS^His^ RNAs using the SHAPE reagent NMIA: Tobacco Mosaic virus (A), Ribgrass Mosaic Virus (B), Hibiscus Latent Fort Pierce virus (C), and Diascia Yellow Mottle virus (D). Reactivity was background subtracted and normalized to flanking 5’ and 3’ normalization hairpins (not depicted, see Supplemental File 2 for sequence details). Shading represents degree of normalized modification according to the insert legend.

### Representative putative new TLS^His^ are histidylated *in vitro*

While all new TLS^His^ conformed to the consensus secondary structure, the observed sequence diversity motivated us to determine if they are indeed substrates for aminoacylation by histidyl-tRNA synthetase (HisRS). We used *in vitro* aminoacylation assays using purified HisRS from *S. cerevisiae* and ^3^H-2,5-*L*-histidine as substrate; incorporation of the ^3^H into the RNA reflects aminoacylation. To test the specificity of this assay, we used a yeast histidine tRNA (tRNA^His^) and a yeast leucine tRNA (tRNA^Leu^) as positive and negative controls, respectively. The yeast tRNA^His^ demonstrated a high level of histidine incorporation while yeast tRNA^Leu^ showed low levels of histidine incorporation (Fig. 3). Consistent with previous reports, the prototypical TLS^His^ from TMV was histidylated at levels matching tRNA^His^ (Fig. 3) (Felden et al. 1994b). We then tested seven representative newly identified putative TLS^His^ RNAs, chosen to contain several variable features including diverse anticodon sequences, discriminator nucleotide identity, AC stem length, and AC bulge size to assay if these features were accommodated by the HisRS. All of these putative TLS^His^ RNAs were histidylated well above the negative controls, indicating they are authentic TLS^His^. Although we did not test every new putative TLS^His^, the fact that all those that were tested were aminoacylated and the robust conservation within the class suggests that most or all of the new examples are likely functional substrates for HisRS.

**Figure 3:**
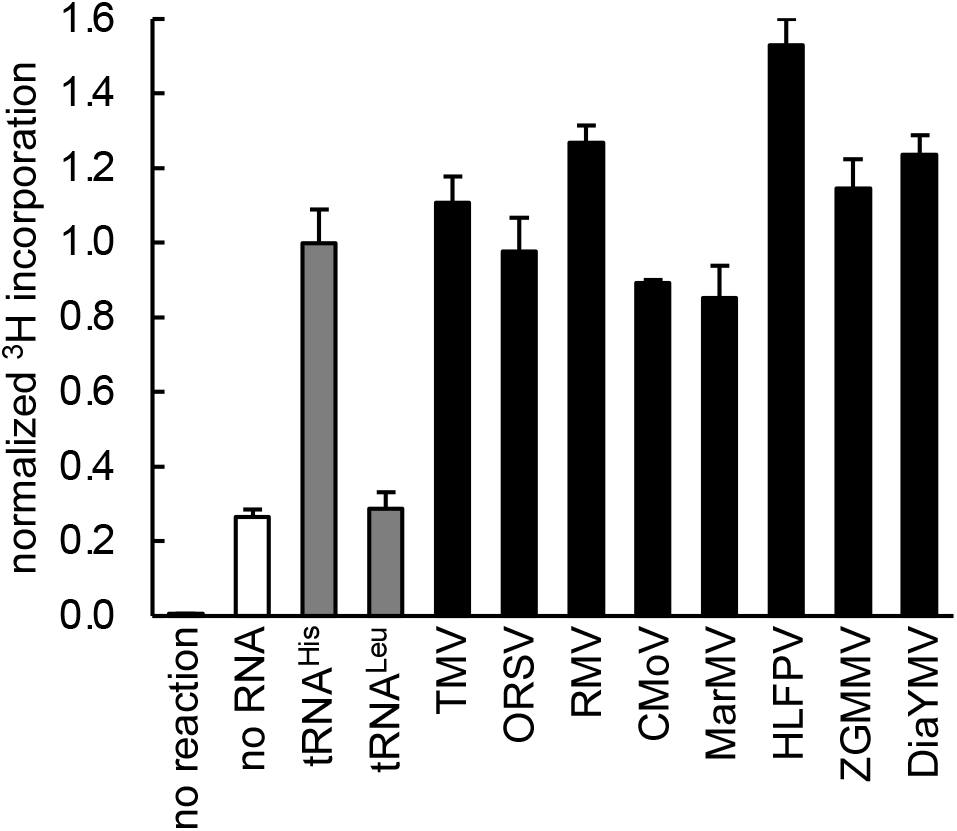
Identified putative viral TLS^His^ sequences are histidylated *in vitro*. ^3^H-L-histidine incorporation of eight representative TLS^His^ RNAs identified through bioinformatic searches. Histidylation of each RNA, as measured by covalent incorporation of ^3^H-L-histidine by histidine tRNA-synthetase (HisRS) at the 3’ adenosine, is normalized to yeast tRNA^His^ RNA. Each reaction was performed in triplicate. Error bars represent one standard error from the mean.

### TLS^His^ aminoacylation bypasses features required in tRNAs

Several important histidine identity nucleotides that facilitate recognition by the HisRS are proposed to be present in TLS^His^ RNAs (Rudinger et al. 1997; Dreher 2010), but we observed that several TLS^His^ which lacked these features maintained the ability to be aminoacylated (Fig. 2 and Supplemental File 1). First, within a canonical tRNA^His^ two primary identity elements are the N_−1_ and N_73_ bases at the end of the acceptor stem (Rudinger et al. 1997; Tian et al. 2015). These are nearly invariant across tRNA^His^ as G_−1_ and A_73_. However, in TLS^His^ the homologous bases are often an A for N_−1_ and C for N_73_ (Fig. 1B), with some variation. Indeed, our results show the discriminator nucleotide is not always essential for histidylation for TLS^His^, as there was no significant decrease in histidine incorporation for DiaYMV, which contains an A at this position in place of the C typical of most TLS^His^ and many tRNA^His^. Notably, in most members of the *Virgaviridae* family the discriminator nucleotide analog N_73_ is a C and in all members of the *Tymoviridae* family it is an A. It appears that rather than reflecting the identity of the canonical discriminator nucleotide, this base reflects the viral RdRp lineage (Deiman et al. 1998; Osman et al. 2000), which appears to also be true in the TLS^Val^ class (Sherlock et al. 2020).

The other major identity element within tRNA^His^ is the AC loop and in the TLS^His^ it is most often a GUG as in canonical tRNA^His^ (Rudinger et al. 1997). However, exceptions to this, namely TMV (GUU), RMV (CGG), and MarMV (AGA) TLSs, were readily aminoacylated (Fig. 3). Of note, the AC loop is not as well conserved as the AC loop of the TLS^Val^ class where the sequence is critical for aminoacylation (Sherlock et al. 2020). Additionally, the TLS^His^ AC loop is predicted to contain five nucleotides with covariation in the terminal base pair in the stem, in contrast with the canonical seven-nucleotide tRNA^His^ AC loop (Rudinger et al. 1997). Finally, in the TLS^His^ class, the AC stem always contains an internal loop or bulge and is significantly longer than the AC stem of the TLS^Val^ class. This difference in stem length likely relates to the three-dimensional structure of the TLS^His^ RNAs though further studies are needed to understand how these differences are resolved in the structure. Perhaps the flexibility in this region affords the structure with dynamic capabilities to switch between different functional states, as proposed for the TLS^Tyr^ (Bonilla et al. 2020); this could explain previous data suggesting conformational heterogeneity in TLS^His^ (Hammond 2009).

### A conserved D-loop/T-loop-like interaction is present in the TLS^His^

Chemical probing showed that in all RNAs, the conserved GG dinucleotides present between PK2 and the AC stem generally had decreased reactivity compared to the adjacent nucleotides and other unpaired regions of RNA, despite no obvious base pairing partners (Fig. 2). Similarly, the T-loop region displayed decreased levels of SHAPE reactivity when compared with other loop and linker regions, though still higher than regions of base pairing. Within the T-loop, the second U was consistently more reactive than the rest of the loop, similar to what is observed in canonical tRNAs (Kladwang et al. 2011), where this T-loop base participates in a long-range interaction with the D-loop forming the distinct elbow of the tRNA (Levitt 1969; Rould et al. 1989; Tian et al. 2015). In canonical tRNAs this elevated reactivity is due to the unique local backbone geometry (Kladwang et al. 2011; Tian et al. 2015). This shared pattern reactivity led us to hypothesize that a similar structure and interaction is present in the TLS^His^ class. Indeed, previous studies proposed this interaction exists in the TMV TLS though it has not been directly tested (Felden et al. 1996).

To test if the TLS^His^ class contains an interaction between the conserved GG dinucleotide and the T-loop, we tested the effect of mutating each of these elements on function and structure. Specifically, we introduced a G➔A mutation in the GG dinucleotide and a C➔U mutation in the T-loop within three representative RNAs: TMV, RMV, and DiaYMV TLS^His^ (Fig. 4A-B and Fig. S2). These mutants were chosen because analogous mutations in tRNAs, G19A and C56U, disrupt the D-loop/T-loop interaction, resulting in decreased aminoacylation (Du and Wang 2003). Specifically, C➔U is located in the T-loop analog and G➔A is in the GG dinucleotide proposed to form a Watson-Crick base-pairing interaction with the T-loop. When these mutant TLS^His^ were tested in our histidine incorporation assay, they exhibited low levels of aminoacylation (Fig. 4C), close to or within error of the tRNA^Leu^ negative control and reactions lacking any RNA. The near total loss of histidine incorporation resulting from these mutations is similar to what has been observed for canonical tRNAs when the T-loop/D-loop interaction is disrupted (Du and Wang 2003).

**Figure 4:**
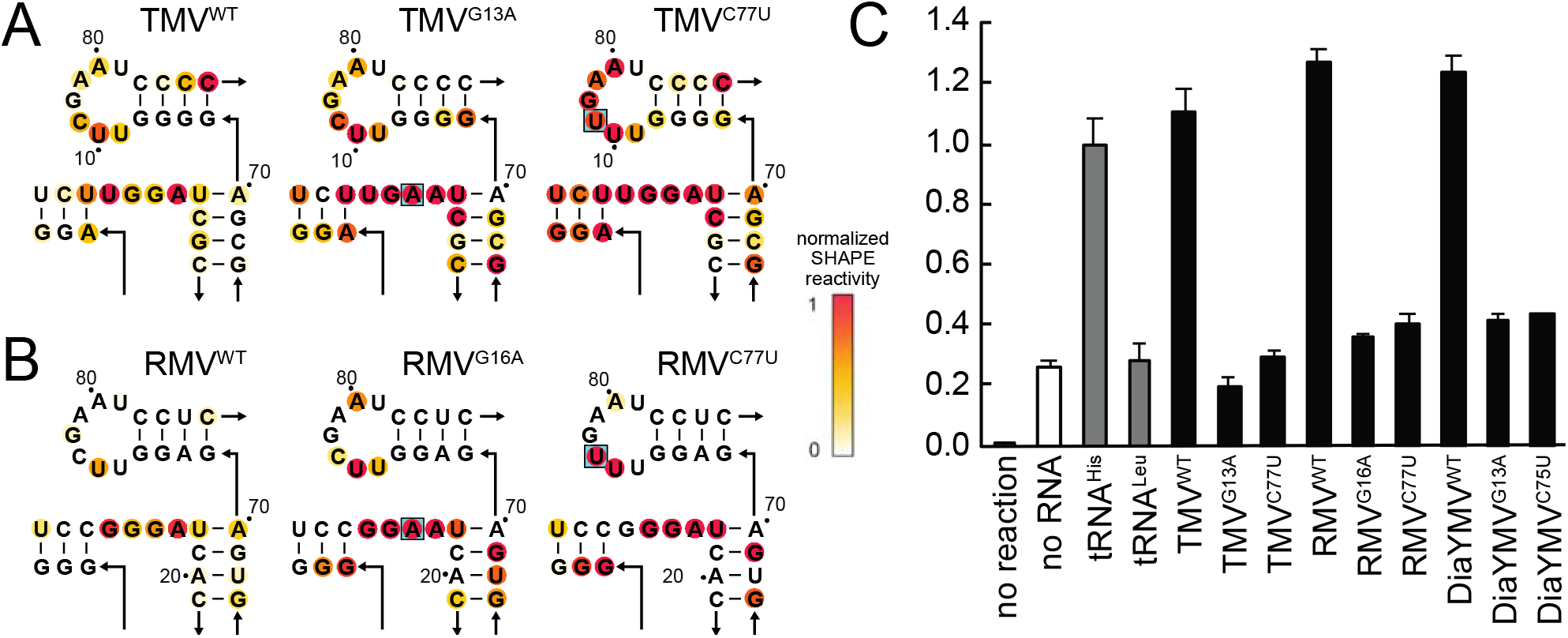
A putative conserved D-loop/T-loop mimic is required for structure and efficient histidylation. **A-B)** SHAPE chemical probing of the wild type TMV (A) and RMV (B) TLSs and two D-loop/T-loop mutants (boxed in blue in figure). Reactivity was background subtracted and normalized to flanking 5’ and 3’ normalization hairpins (not shown). **C)** ^3^H-L-histidine incorporation of three representative TLS^His^ RNAs; TMV, RMV, and DiaYMV, each with the wild type sequence as well as two D-loop/T-loop mutants. Histidylation of each RNA, as measured by covalent incorporation of ^3^H-L-histidine by histidine tRNA-synthetase (HisRS), is normalized to yeast tRNA^His^. Each reaction was performed in triplicate. Error bars represent one standard error form the mean.

The aminoacylation assays reveal the functional importance of these specific nucleotides, but do not show if they interact, therefore we used chemical probing to determine if mutation to either the T-loop or GG dinucleotide induced changes in other parts of the RNA. SHAPE probing of the C➔U RNAs revealed substantial increases in reactivity in the T-loop, consistent with destabilization of the loop’s structure. In addition, the reactivity of the GG dinucleotides also increased, while changes in the rest of the RNA were minimal (Fig. S2). Thus, disruption of the T-loop in these TLS^His^ causes discrete structural changes in the GG dinucleotide, consistent with a long-range interaction between these elements. The G➔A mutation (in the GG dinucleotide) induced increased reactivity in the GG dinucleotide and some subtle increases in the T-loop (Fig. 4A-B). This is consistent with the proposed interaction and likely reflects the fact that the T-loop comprises a preformed structural motif whose structure is less dependent on tertiary interactions (Chan et al. 2013).

Our data suggest that the GG dinucleotide between PK2 and the AC stem of TLS^His^ acts analogously to the D-loop in tRNAs, making a specific long-range contact to the T-loop as previously proposed (Felden et al. 1996). We speculate this interaction consists of a Watson-Crick base pair between the first G of the dinucleotide, G_12_ in our TMV construct, and the third base in the T-loop, C_78_, as well as the reverse Hoogsteen base pair between G_13_ and U_76_, as seen in tRNAs. In tRNAs this interaction is critical to create the functional global fold, and this is likely true of the TLS^His^ although high-resolution structural information will be needed to test this hypothesis. If true, the GG dinucleotide ‘D-loop’/T-loop interaction mimic in TLS^His^ likely plays a similar structural and functional role as the canonical D-loop/T-loop interaction in tRNAs, positioning the AC stem and acceptor pseudoknot such that the HisRS can productively recognize these important identity elements.

### Comparisons between the TLS^His^ and other TLS classes

In some ways, characteristics of the TLS^His^ class lay in the middle of the three known TLS classes. For example, the identity and size of the AC loop is strictly conserved for TLS^Val^ and examples that lack an authentic valine anticodon are not aminoacylated by valyl-tRNA synthetase (Sherlock et al. 2020). In contrast, the AC loop is not conserved for the TLS^Tyr^ class, to the point where there are disagreements in its identity (Perret et al. 1989; Felden et al. 1994a; Bonilla et al. 2020). While many TLS^His^ representatives contain a GUG histidine AC, there are many examples that do not, and the AC loop length for TLS^His^ is typically only five instead of seven nucleotides. Despite this, all are still aminoacylated by HisRS. Additionally, the TLS^His^ class contains more secondary structure elements than TLS^Val^ and fewer than TLS^Tyr^ and also falls in the middle in average length. Perhaps this intermediate status could help elucidate the evolutionary relationship and trajectory of the different TLS classes. While our current studies do not address these evolutionary relationships, it is noteworthy that the TLS^His^ and TLS^Val^ structures have enough similarity, especially in the 3′ regions, that bioinformatic searches based on the TLS^His^ class can find TLS^Val^ and *vice versa,* albeit mostly at E values above the inclusion threshold (Sherlock et al. 2020). It also remains to be determined how the TLSs achieve specificity for a particular amino acid given the observed diversity in identity elements. It is intriguing to speculate that other classes of TLSs exist that are structurally unique compared to the three known classes. There is certainly precedent to support this, given the constantly expanding list of classes of riboswitches, ribozymes, and xrRNAs as well as the existence of other tRNA-mimicking structures such as TMERs, tmRNAs, and mascRNAs.

## Concluding Remarks

In this study we expanded the list of known TLS^His^, confirmed their shared secondary structure, shown they are histidylated *in vitro*, and validated a proposed conserved D-loop/T-loop interaction analog. These discoveries present new questions regarding how this class of TLSs achieves specific interactions required for recognition by both host and viral proteins while demonstrating significant sequence and secondary structure variation from canonical tRNAs. While the experiments and analyses herein build on and confirm previous observations and hypotheses regarding TLS^His^, high-resolution structural information on TLS^His^ RNAs will be necessary to address lingering questions regarding their structure and function. For example, in this class the AC-arm is particularly long compared to canonical tRNA^His^ and always has an internal loop. How this additional length is accommodated, if the loop affects orientation of the anticodon relative to a bound HisRS and if these elements provide conformational dynamics or reconfigurations remains to be understood. Overall, this study moves us closer to understanding the molecular interactions underlying tRNA mimicry for the TLS^His^ class in terms of structure, while the overall topology, intermolecular interactions with host and viral proteins, and ultimately the function of these RNA elements during infection, remains to be elucidated.

## MATERIALS AND METHODS

### TLS^His^ bioinformatic searches and consensus model generation

An alignment of four known histidine-accepting tRNA like structures (TLS^His^) (Rfam ID: RF01077) was obtained from the Rfam database (Kalvari et al. 2018) and extended to include the entire TLS, as the existing alignment ended prior to the T-loop and PK1. Using Infernal version 1.1 (Nawrocki and Eddy 2013) a database consisting of all +ssRNA virus sequences deposited in the National Center for Biotechnology Information (NCBI, retrieved 01/22/2019) was queried to identify additional instances of this motif. Sequences identified by Infernal were added to the initial four sequences to generate an updated covariance model for subsequent iterative searching. Only sequences below the Infernal E-value threshold of 0.05 were considered. Duplicate sequences were removed yielding 157 sequences from 35 unique viruses. These resulting sequences were used to generate a consensus sequence and secondary structure model including an analysis of covariance using RNA Covariation Above Phylogenetic Expectation (R-scape) (Rivas et al. 2017, Rivas et al. 2020) then rendered in R2R (Weinberg and Breaker 2011).

### Expression of *Saccharomyces cerevisiae* HisRS

The DNA sequence encoding HisRS enzyme from *S. cerevisiae* (GenBank: AJW07132.1) was purchased as a dsDNA gBlock (IDT) and cloned into a pET15b(+) vector containing an in-frame N-terminal hexahistidine affinity tag. The protein was recombinantly expressed in BL21 (DE3) cells. Cells were grown in LB to an OD_600_ of 0.3, then protein expression was induced using 250 μM isopropyl β-D-1-thiogalactopyranoside (IPTG) overnight at 18°C. Pelleted cells were resuspended in lysis buffer containing 20 mM Tris-HCl (pH 7.0), 500 mM NaCl, 2 mM β-mercaptoethanol (BME), 10% (v/v) glycerol, and cOmplete EDTA-free protease inhibitor cocktail tablets (Roche). Cell lysate was then sonicated on ice for 2 minutes of: 20 seconds on, 40 seconds off at 75 W. Cell lysate was clarified by centrifugation at 30,000xg for 30 minutes at 4°C. The soluble fraction was purified by nickel affinity chromatography in a buffer containing 150 mM NaCl, 20 mM Tris (pH 7.0), 200 mM Imidazole, 10% glycerol, and 2 mM β-mercaptoethanol. The protein was exchanged into a storage buffer containing 50 mM Tris-HCl (pH 8.0), 100 mM NaCl, 5 mM MgCl_2_, and 5% glycerol using a spin concentrator (Amicon) and stored at 0.3 mg mL^−1^ at −80°C with working stocks stored at −20°C.

### *In vitro* RNA transcription

DNA templates were ordered as gBlock DNA fragments (IDT) and cloned into pUC19. 200 μL PCR reactions using primers containing an upstream T7 promoter were carried out to generate dsDNA templates for transcription. Typical PCR conditions: 100 ng plasmid DNA, 0.5 μM forward and reverse DNA primers (see Supplemental File 2), 500 μM dNTPs, 25 mM TAPS-HCl (pH 9.3), 50 mM KCl, 2 mM MgCl_2_, 1 mM β-mercaptoethanol, and Phusion DNA polymerase (New England BioLabs). Templates for RNA used in aminoacylation assays were amplified using reverse primers containing two 5′- terminal 2′-*O*-methyl modified bases to ensure the correct 3′ end of the RNA. dsDNA amplification was confirmed by 1.5% agarose gel electrophoresis. Transcriptions were performed in 1 mL volume using 200 μL of PCR product. Transcription conditions: ~ 0.1 μM template DNA, 10 mM NTPs, 75 mM MgCl_2_, 30 mM Tris (pH 8.0), 10 mM DTT, 0.1% spermidine, 0.1% Triton X-100, and T7 RNA polymerase. Reactions were incubated at 37°C overnight. After transcription, insoluble inorganic pyrophosphate was removed by centrifugation at 5000xg for 5 minutes then the RNA-containing supernatant was ethanol precipitated with 3 volumes of 100% ethanol at −80°C for a minimum of 1 hour then centrifuged at 21000xg for 30 minutes at 4°C to pellet the RNA and the ethanolic fraction was decanted. The RNA was resuspended in 9 M urea loading buffer then purified by denaturing 10% PAGE. Bands were visualized by UV shadowing then excised. Bands were then crush-soaked in diethylpyrocarbonate-treated (DEPC) milli-Q water at 4°C overnight. The RNA-containing supernatant was then concentrated using spin concentrators (Amicon) to the appropriate concentration. RNAs were stored at −80°C with working stocks stored at −20°C.

### *In vitro* chemical probing of RNAs

Structure probing experiments using the SHAPE reagent NMIA were performed as described previously (Cordero et al. 2014). Briefly, 240 μM RNAs were refolded by heating to 90°C for 5 minutes, cooled to ambient temperature, then incubated at ambient temperature with MgCl_2_ for 20 minutes. Subsequently, the refolded RNA was modified by incubating with NMIA for 15 minutes at ambient temperature. NMIA modification conditions: 120 nM RNA, 6 mg/mL NMIA or DMSO, 50 mM HEPES-KOH (pH 8.0), 10 mM MgCl_2_, 3 nM 6-fluorescein amidite 5′-labeled FAM-RT primer (see Supplemental File 2). Modification was quenched by the addition of NaCl to 500 mM, Na-MES buffer (pH 6.0) to 50 mM, and oligo-dT magnetic beads (Invitrogen Poly(A)Purist MAG Kit). Modified RNAs were recovered using the magnetic beads then washed twice with 70% ethanol then resuspended in water. Reverse transcription was carried out using SuperScript III (Invitrogen) at 48°C for 1 hour per the manufacturer’s instructions. The RNA was then degraded by the addition of NaOH to 200 mM and heating to 90°C for 5 minutes. An acid-quench solution (final concentration: 250 mM NaOAc (pH 5.2), 250 mM HCl, 500 mM NaCl) was added and DNA was recovered using the magnetic beads. The DNA was washed twice with 70% ethanol and eluted in GeneScan 350 ROX Dye Size Standard (ThermoFisher) containing HiDi formamide solution (ThermoFisher). 5′-FAM labeled reverse-strand DNA products were resolved by capillary electrophoresis using an Applied Biosystems 3500 XL instrument. Data workup was performed using the HiTrace RiboKit (https://ribokit.github.io/HiTRACE/) (Yoon et al. 2011; Kim et al. 2013; Kladwang et al. 2014; Lee et al. 2015) in MatLab (MathWorks), and figures were rendered using RiboPaint (https://ribokit.github.io/RiboPaint/) in MatLab then labeled in Adobe Illustrator. SHAPE reactivity was superimposed on the predicted secondary structure and used to make adjustments to the secondary structure.

### *In vitro* aminoacylation assays

Aminoacylation constructs were refolded by heating to 90°C for 5 minutes then cooling to ambient temperature, then incubated with 10 mM MgCl_2_ for 20 minutes. Aminoacylation reactions were setup as follows; 1 μL of 1 μM RNA or water, 1 μL of freshly prepared aminoacylation buffer (10x: 200 mM HEPES-KOH (pH 7.5), 20 mM ATP, 300 mM KCl, 50 mM MgCl_2_, 50 mM DTT), 1 μL of ^3^H-2,5-L-histidine, 6 μL of DEPC-treated water, and 1 μL of HisRS (3 μM). Aminoacylation reactions were incubated at 30°C for 2 hours. Reactions were quenched with 100 μL of wash buffer (20 mM Bis-Tris (pH 6.5), 10 mM NaCl, 1 mM MgCl_2_) with trace xylene cyanol for visualization. Quenched reactions were immediately loaded onto a vacuum filter blotting apparatus. Filter stack in order from top to bottom: 0.45 μM Tuffryn membrane filter paper (PALL Life Sciences), HyBond positively charged membrane (GE Healthcare), thick filter paper (Bio-Rad gel dryer filter paper). Prior to filter blotting apparatus assembly, each layer was equilibrated in wash buffer. After application of the reaction solution, each blot was immediately washed with 3×300 μL of wash buffer containing trace xylene cyanol. The filters were subsequently dried and the blots from the HyBond membrane were excised and measured for ^3^H incorporation by liquid scintillation counter (Perkin-Elmer Tri-Carb 2910 TR). Data processing was performed in Excel.

## Supporting information

Sequence alignment

Accession numbers, sequences, and raw data

## ACKNOWLEDGEMENTS

The authors thank the members of the Kieft Lab for thoughtful discussions and David Costantino for critical reading of this manuscript. This work was supported by NIH grant R35GM118070 to JSK. MES is a Jane Coffin Childs Postdoctoral Fellow.

