## Supplemental Figures for "An Expanded Class of Histidine-Accepting Viral tRNA-like Structures"

**Table of Contents:**

Figure S1: Chemical probing data for diverse $TLS^{His}$
Figure S2: Chemical probing data for D-loop/T-loop mimic mutants
Figure S3: Additional D-loop/T-loop mutant $^{3}H$-incorporation assays

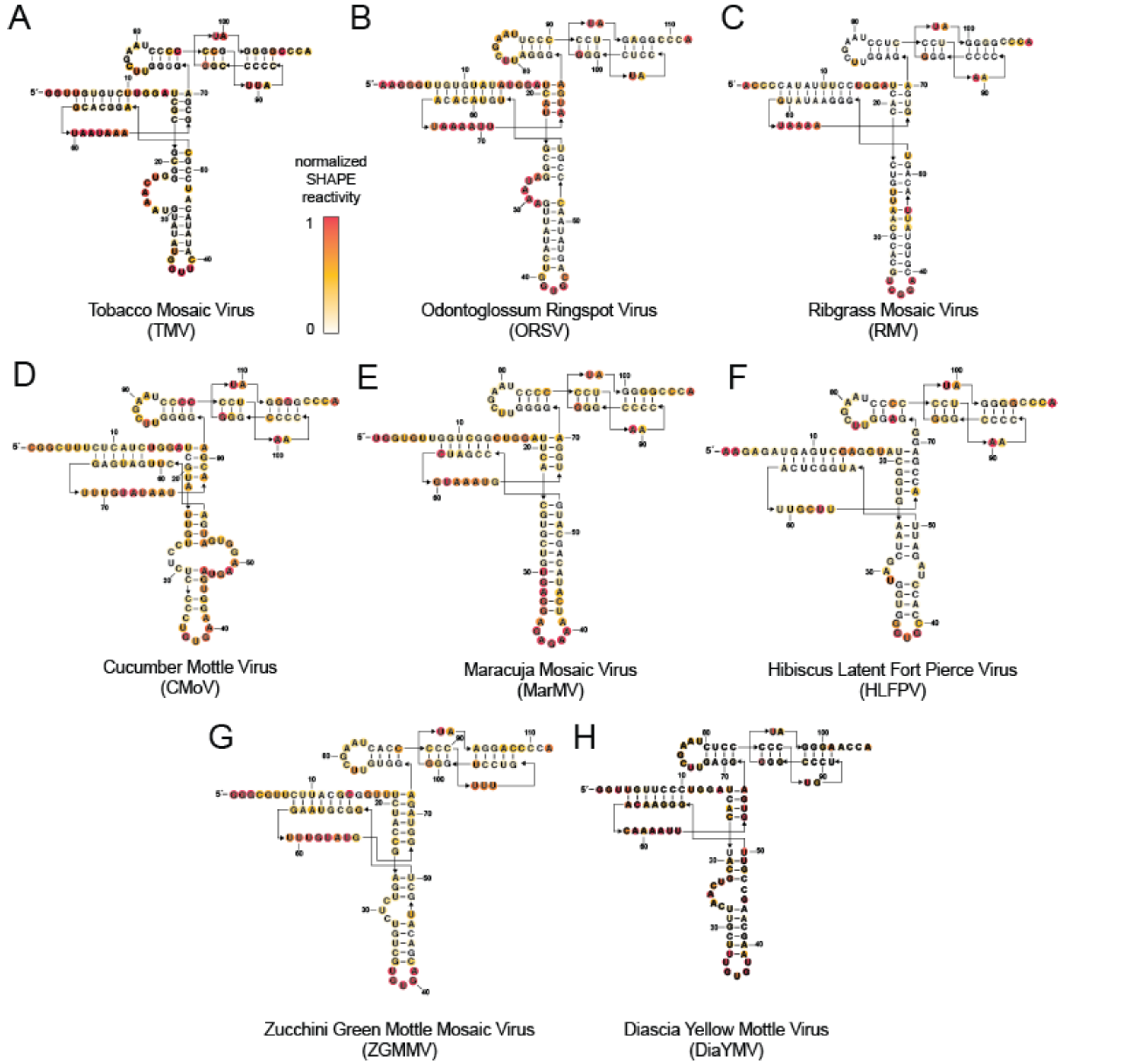


**Figure S1. Chemical probing data for diverse TLS$^{His}$. A-H)** Chemical probing of all representative TLS$^{His}$ RNAs using the SHAPE reagent NMIA: Tobacco Mosaic virus (A), Odontoglossum Ringspot Virus (B), Ribgrass Mosaic Virus (C), Cucumber Mottle Virus (D), Maracuja Mosaic Virus (E), Hibiscus Latent Fort Pierce Virus (F), Zucchini Green Mottle Mosaic Virus (G), and Diascia Yellow Mottle Virus (H). Reactivity was background subtracted and normalized to flanking 5' and 3' normalization hairpins (not depicted, see Supplemental File 2 for sequence details).

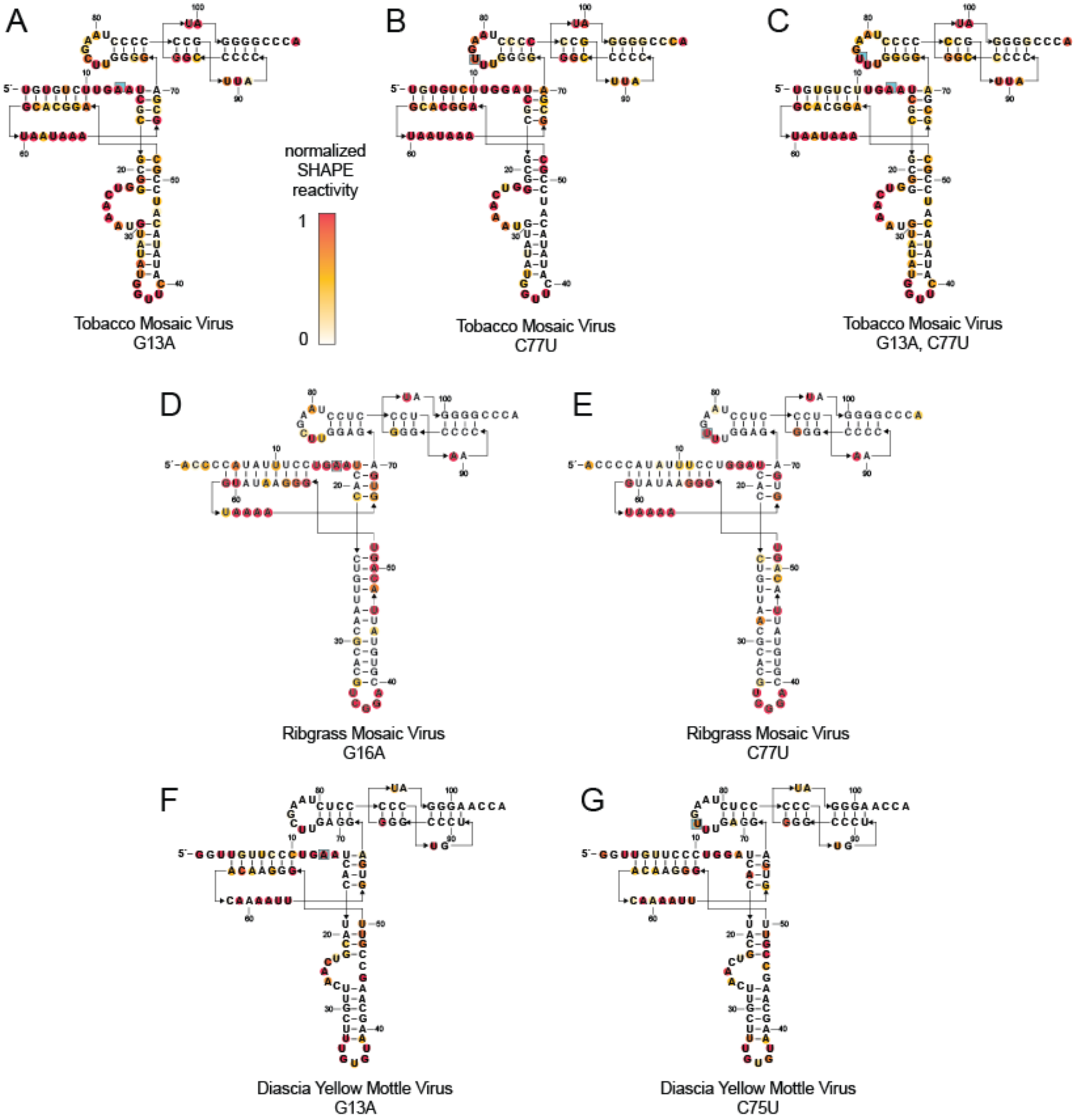


**Figure S2. Chemical probing data for D-loop/T-loop mimic mutants. A-G)** Chemical probing of all mutant D-loop/T-loop mimic TLS$^{His}$ RNAs using the SHAPE reagent NMIA: Tobacco Mosaic virus G13A (A), Tobacco Mosaic virus C77U (B), Tobacco Mosaic virus G13A, C77U (C), Ribgrass Mosaic Virus G16A (D), Ribgrass Mosaic Virus C77U (E), Diascia Yellow Mottle Virus G13A (F), and Diascia Yellow Mottle Virus C75U (G). Reactivity was background subtracted and normalized to flanking 5’ and 3’ normalization hairpins (not depicted, see Supplemental File 2 for sequence details).

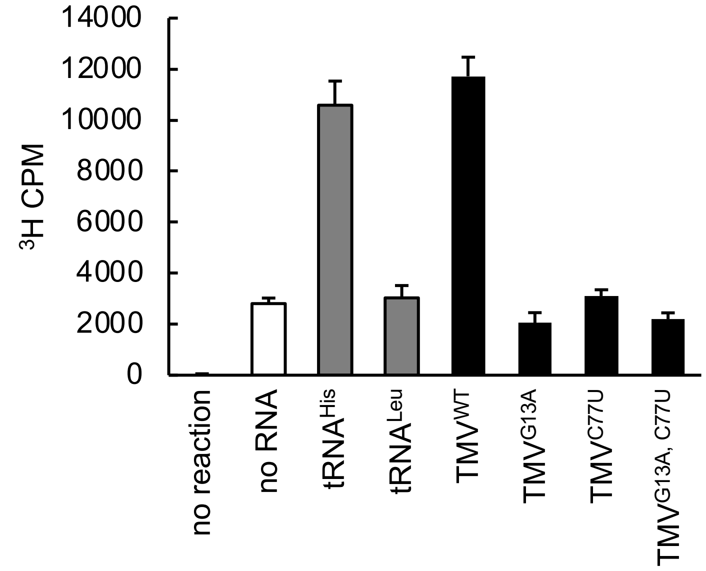


**Figure S3. Additional D-loop/T-loop mutant $^3$H-incorporation assays.** $^3$H-L-histidine incorporation of four TMV TLS$^{His}$ RNAs: TMV$^{WT}$, TMV$^{G13A}$, TMV$^{C77U}$, and TMV$^{G13A,\ C77U}$. Histidylation of each RNA, as measured by covalent incorporation of $^3$H-L-histidine by histidine tRNA-synthetase (HisRS), is normalized to yeast tRNA$^{His}$. Each reaction was performed in triplicate. Error bars represent one standard error from the mean.
